# Cell rearrangement progression along the apical-basal axis is linked with 3D epithelial tissue structure

**DOI:** 10.1101/2024.04.29.591620

**Authors:** Erika M. Kusaka, Sassan Ostvar, Xun Wang, Xiaoyun Liu, Karen E. Kasza

## Abstract

Epithelial tissues undergo extensive structural remodeling during embryonic development. Tissue remodeling is often enabled by oriented cell rearrangements that are linked with patterns of mechanical stress in the tissue and with tissue mechanical properties. Cell rearrangements and their links to tissue structure have largely been studied at the apical side of tissues at the level of adherens junctions. Less is known about the involvement of basolateral domains in cell rearrangements. Here we use live confocal imaging to quantify cell rearrangements, cell packing structure, and cell morphology in 3D in the converging and extending *Drosophila* germband epithelium. We report gradients in cell shapes and tissue structure along the apical-basal axis of the germband, suggesting that the apical and basolateral domains display distinct behaviors. Cell rearrangements initiate at apical as well as basolateral positions, with initiation frequencies also displaying a gradient along the apical-basal axis. Following initiation, rearrangements propagate across the apical-basal axis and lateral cell contacts remodel; these events involve scutoids and other complex 3D cell shapes as intermediate states. These findings uncover novel aspects of the cell rearrangements that drive dynamic remodeling of epithelia and reveal links between rearrangements and gradients in tissue structure along the apical-basal axis.

---

Epithelial tissue sheets dynamically remodel and flow during development and in response to injury (1, 2). A primary mechanism driving epithelial remodeling in many contexts is local cell rearrangements that allow remodeling of cell-cell contacts without opening gaps in the tissue (3–5). Significant research has focused on the proteins organizing junctional remodeling during cell rearrangements at the apical side of tissues (5–12), and how apical cell packing structure influences the solid-fluid tissue mechanical properties that resist tissue flow (13–19). Less attention has been focused on the involvement of basolateral domains in cell rearrangements, although there is increasing evidence that basal protrusive activity contributes to cell rearrangements in many contexts in both epithelial and mesenchymal tissues (3, 20–23). It remains poorly understood how epithelial cell rearrangements occur along the full apical-basal axis in the three-dimensional context of epithelia. Coupling between apical and basal sides of the tissue during cell rearrangements may be especially crucial in tall epithelia in which the apical and basal sides of the tissue are physically distant.

Recent studies have revealed that complex 3D cell shapes naturally arise in static epithelial tissue sheets that are curved or tilted. In these contexts, some columnar epithelial cells no longer take on the shapes of simple frusta and instead have different lateral cell neighbor contacts along the apical-basal axis in order to accommodate overall tissue shape (24–28). This results in static “apical-basal rearrangements” in which cells exchange neighbors along the apical-basal axis instead of along the time axis. These structures are associated with cell geometries known as *scutoids* (24, 29). Transient apical-basal rearrangements and scutoids have recently been reported in proliferating epithelia in which new cells are generated and must be accommodated (28). However, it is not known what 3D cell geometries might be involved in dynamic cell rearrangements in remodeling or flowing tissues, even in the context of relatively flat, non-proliferating tissue sheets.

Here, we combine live confocal imaging, quantitative image analysis, and 3D reconstructions of remodeling cell-cell contacts to study how cell rearrangements proceed along the apical-basal axis over time in the converging and extending *Drosophila* germband epithelium. First, we show that cell shapes and tissue structure vary along the apical-basal axis of the tissue, suggesting varying behavior along this tissue axis. Next, we find that cell rearrangements can initiate at any depth along the apical-basal axis, although initiation is most frequent near the apical side of the tissue; rearrangements then propagate across the entire apical-basal axis of the tissue to produce complete cell neighbor exchanges. Notably, the apical-basal gradient in cell rearrangement initiation frequencies mirrors the gradient in tissue structure along this axis. Finally, we explore the 3D geometries of cells that participate in dynamic cell rearrangements in the germband and find that scutoids and other complex geometries represent transient intermediate states in rearrangements as lateral cell-cell contacts remodel, revealing novel features of cell rearrangements that mediate epithelial tissue shape changes.

## Cell shapes and tissue structure vary along the apical-basal axis of the *Drosophila* germband epithelium

We began by exploring how tissue structure varies along the apical-basal axis of the *Drosophila* germband epithelium. The germband converges and extends to elongate the head-to-tail body axis of the embryo in a process known as germband extension (GBE) (Fig. 1A) (6). This rapid tissue flow is mediated by oriented cell rearrangements that are thought to be driven by planar polarized patterns of actomyosin contractility at adherens junctions near the apical side of the tissue (7–9), although cell activities at the basal side have also been reported (21). Recently, our group and others showed that the onset of cell rearrangements can be predicted by cell shapes at apical cross-sections of the tissue, which can be understood as a transition from solid-like to more fluid-like tissue behavior (14, 15, 19). However, it is not known whether cell shapes vary along the apical-basal axis of the ≈30 micron tall columnar germband epithelium or how any variations be might be linked with tissue remodeling.

**Fig. 1.**
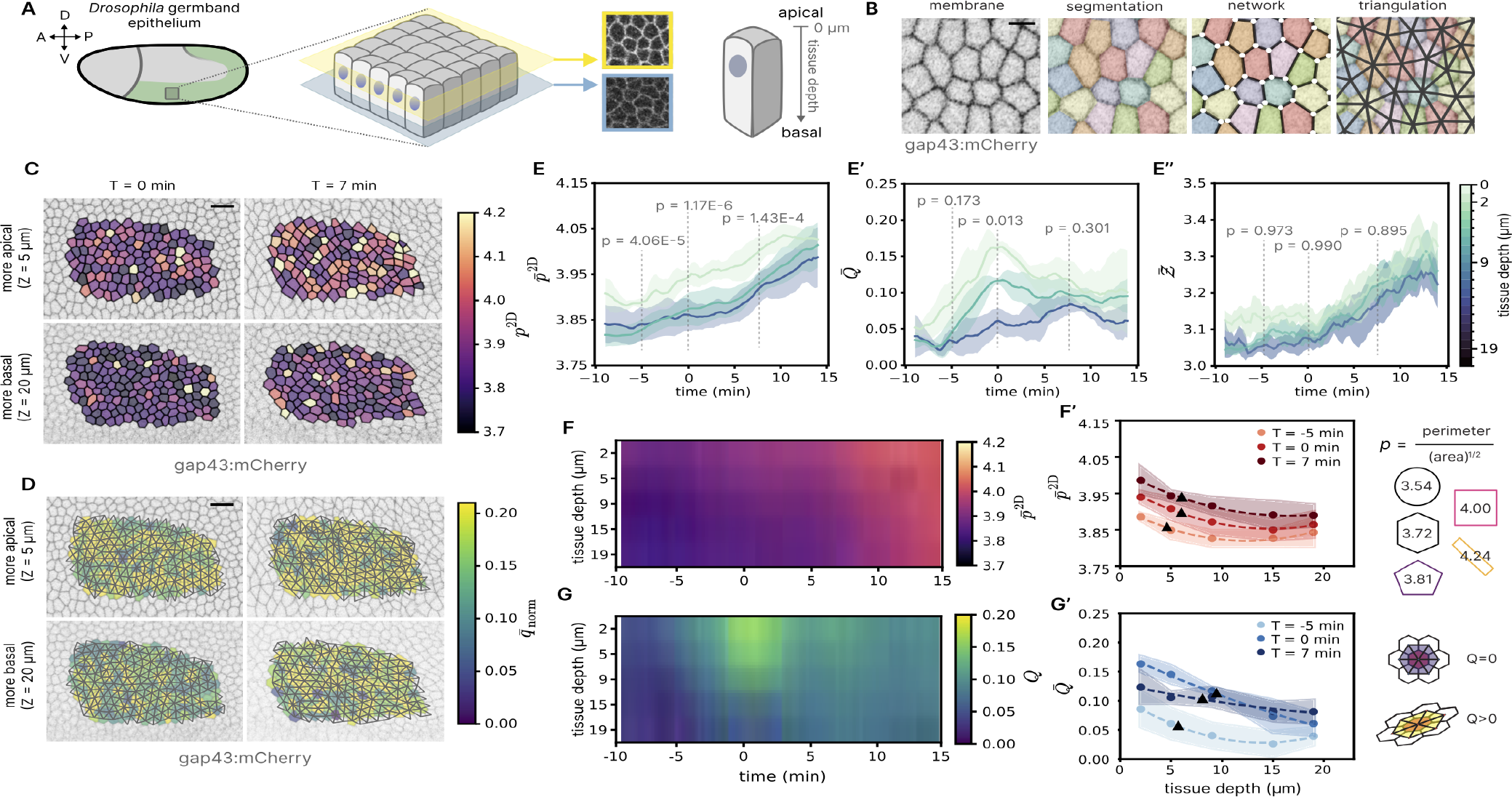
Cell shapes and tissue structure vary along the apical-basal axis in the *Drosophila* germband epithelium. **(A)** Schematic of the *Drosophila* germband epithelium during germband extension (GBE) (left). Schematic of the germband epithelial tissue with example confocal images of cell shapes at apical and basolateral positions in the germband. Schematic of apical-basal axis mapped onto a cell with z=0 set at the apical surface. **(B)** Cell membranes labeled with gap43:mCherry are visualized by confocal microscopy. At each z-position along the apical-basal axis, cells are segmented and represented as polygons that tile the plane to quantify cell shapes and vertex coordination number. The polygon network is triangulated for quantification of cell shape alignment. Scale bar, 5 microns. **(C)** The cell shape index from confocal z-slices, *p*^2D^, for individual cells at apical (top) and basolateral (bottom) z-positions in the tissue at the onset of GBE (T=0 min, left) and 7 min after the onset of GBE (right). Scale bar, 15 microns. **(D)** Triangulation used to calculate the cell shape alignment index, *Q*. The color of each cell represents the local tissue alignment quantified by the average norm of the triangle nematic directors (*q*) for triangles that have a vertex at the cell centroid. Scale bar, 15 microns. **(E)** Time-series of the average cell shape index, 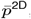, **(E’)** the cell shape alignment index, *Q*, and **(E”)** the vertex coordination number, *Ƶ*, at 3 z-positions along the apical-basal axis of the tissue. For (E), (E’), and (E”), data are the average of N=3 embryos, and shaded regions represent the standard deviation (SD) between embryos. p-values at time points -5, 0, and 7 min (Kruskal-Wallis test). **(F)** Heat map of 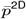 along the apical-basal axis over time. Data represent the average of N=3 embryos. **(F’)** 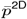 (*z*) at three time points. For each time point, the data is fit to a polynomial curve (dashed line) to determine the characteristic lengthscale at which 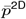 (*z*) falls to half of its maximum value (black triangles). Data are the average of N=3 embryos, and shaded regions represent the SD between embryos. Right: Schematic of the cell shape index with example shapes. **(G)** Heatmap of *Q* along the apical-basal axis over time. Data represent average of N=3 embryos. **(G’)** Average *Q* along the apical-basal axis at three time points. For each time point, the data are fit to a polynomial curve to determine the characteristic lengthscales (black triangles). Data are the average of N=3 embryos, and shaded regions represent the SD between embryos. Right: Schematic of *Q* with examples of low and high alignment.

We acquired confocal timelapse movies of the germband of wild-type embryos expressing a cell membrane marker with z-slices taken at 1 micron intervals and spanning the entire tissue (Fig. 1A). We segmented the images at z-slices with sufficient image quality (typically z = 2-21 microns)(Fig. 1B) and quantified cross-sectional cell shapes in each z-slice using the average cell shape index, 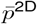, which characterizes whether cells have long or short contacts with their neighbors in the 2D cross section (Fig. 1B) (13, 14, 16, 30). Consistent with prior reports, 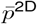 increases over time at the apical side of the tissue (Fig. 1C, E, F) (19). This is accompanied by an increase in vertex coordination number, Ƶ, over time (Fig. 1 E”), which reflects an increase in high-order vertices that form transiently as intermediates during cell rearrangements as the tissue remodels. The increase in 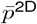 over time is observed in all z-slices, but we also find that 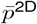 varies along the apical-basal axis (Fig. 1C, E, F). 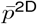 is highest at the apical side of the tissue and then decreases in a gradient towards the basal side (Fig. 1F). The gradient arises from 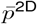 increasing earlier at apical compared to basolateral positions around the onset of GBE (T=0 min) (Fig. 1E, F). The gradient becomes less pronounced by around T = 10 min as 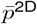 increases at more basolateral positions (Fig. 1F).

Because the apical side of the germband is anisotropic in the plane of the tissue (19), we also quantified anisotropy using the cell shape alignment index, *Q*, derived from a triangulation of cell centroids (Fig. 1B, D) (31). Consistent with prior work, at the apical side of the tissue there is a peak in *Q* around T=0 (Fig. 1E’, G), due to stretching of the germband by the invaginating ventral furrow (19, 32, 33). We again find that alignment is highest at the most apical z-slices (Fig. 1D, E’, G), suggesting that the stretch from ventral furrow invagination most strongly affects the apical side of the tissue. Therefore, similar to 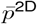, *Q* exhibits an apical-basal gradient.

We wondered how the observed apical-basal gradients in planar cell packing structure relate to epithelial cell organization. To estimate a characteristic lengthscale for the apical-basal gradients, we calculated the tissue depth at which 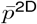 and *Q* reached half their maximum apical values. We found the characteristic lengthscale is ∼6 microns for 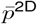 and ∼8 microns for *Q* for the analyzed tissue depth (Fig. 1F’, G’). Both vary weakly over time with some embryo-to-embryo variability (Fig. 1F’ G’). These lengthscales correspond to tissue depths just below the level of apical adherens junctions. The appearance of the apical-basal gradients in cell packing structure correlates with the onset of anisotropic actomyosin-generated stresses at adherens junctions (7, 8, 10, 34–36), suggesting that these gradients may arise from propagation of apical stress patterns along the apical-basal axis.

In 2D vertex models of epithelial tissues (37), preferred cell shapes and tissue structure are predictive of a tissue’s propensity for cell rearrangements (14–17, 19, 38–40). A critical value of 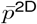 distinguishes between rigid, solid-like and floppy, fluid-like behavior, with higher values of 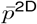 associated with more fluid-like behavior and lower barriers to cell rearrangement in isotropic tissues (14–17, 19). This might suggest more fluid-like behavior at the apical side of the germband around GBE onset. However, it’s unclear how these 2D predictions are related to the 3D picture of epithelia that we present here. Moreover, in anisotropic tissues like the germband, cell shape alignment *Q* increases the critical 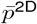 for the rigidity transition (19). Therefore, the apical-basal gradient in 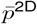 does not necessarily indicate a strong apical-basal gradient in rigidity within the vertex model framework, but the gradient is likely linked to how tissue remodeling proceeds along the apical-basal axis.

Taken together, these results demonstrate that there are structural gradients along the apical-basal axis of the germband that evolve over time during convergent extension, suggesting varying environments along this axis that could potentially be linked to the progression of cell-cell contact remodeling during cell rearrangements.

### Cell rearrangements initiate at various tissue depths and then propagate across the entire apical-basal axis of the tissue

We noticed in our structural measurements that 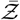 is somewhat higher at more apical z-slices around the onset of GBE (Fig. 1E”). Although this effect is subtle, it hinted that cell rearrangements might initiate at the apical side of the tissue, near actomyosin enrichment at adherens junctions, around the onset of GBE and then propagate across the tissue. To investigate this possibility, we first analyzed a set of manually identified cell rearrangements over time and along the apical-basal axis (Fig. 2A), focusing on 4-cell rearrangements called T1 processes.

**Fig. 2.**
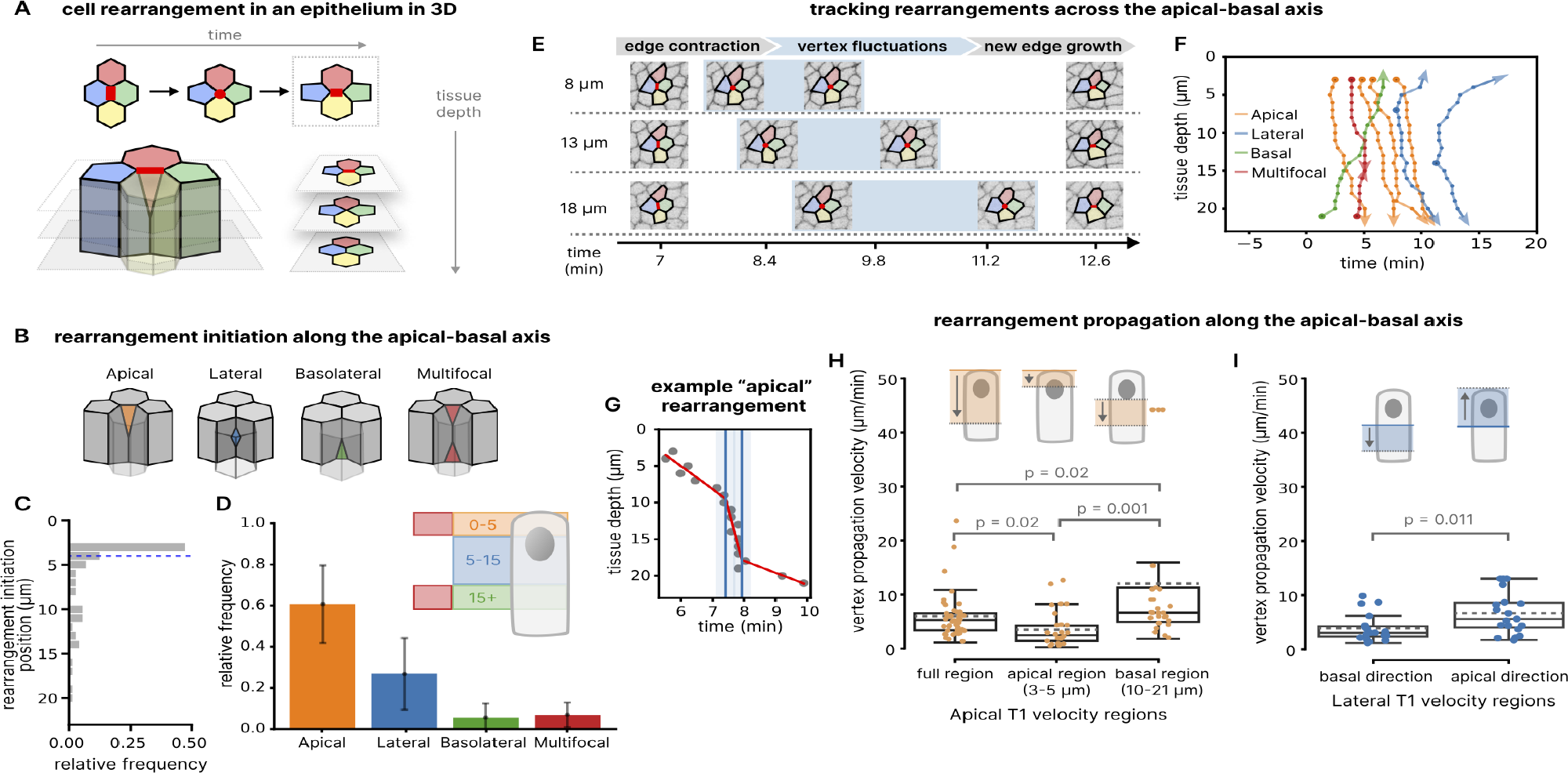
Cell rearrangements initiate at various tissue depths and then propagate along the apical-basal axis. **(A)** Schematic of a cell rearrangement involving four cells (T1 process) that initiates at the apical side of the tissue, highlighting how cell contacts change in time and space. **(B)** Schematics of rearrangements that initiate at different positions along the apical-basal axis. **(C)** Histogram of rearrangement initiation locations along the apical-basal axis (T1 processes only). Position of half the cumulative sum of rearrangements is represented by the blue dashed line. n=73 rearrangements were pooled from N=3 embryos. **(D)** Frequency of each type of rearrangement. Mean and SD between N=3 embryos. Inset: Rearrangement initiation was classified as apical for 0-5 microns, lateral for 5-15 microns, basolateral for 15+ microns. Rearrangement initiation was classified as multifocal if there were both apical and basolateral sites of initiation. **(E-F)** Manual tracking of cell rearrangement initiation and propagation along the apical-basal axis in confocal z-stacks. **(E)** Example of a rearrangement that initiates at the apical side. Images highlighting the four cells in the rearrangement at three z-positions and various time points. Vertices (red circles), changing cell contacts (red lines). Blue regions, time points over which the vertex fluctuates before resolving to form a new contact. **(F)** Example vertex trajectories along the apical-basal axis over time for different types of rearrangements. Vertex initiation points (circles) and direction of vertex propagation (arrow heads) are indicated. **(G)** Example of a vertex trajectory with a piece-wise linear regression analysis (red lines) used to calculate vertex propagation velocities. **(H)** Vertex propagation velocity magnitudes for apically initiated rearrangements. Velocity varies along the apical-basal axis. Median, solid line. Mean, dashed line. Friedman test, p=6.25E-7; p-values from a post-hoc Nemenyi test shown on plot. **(I)** Vertex propagation velocity magnitudes for laterally initiated rearrangements. Velocities are separated into propagation in the apical and basal directions. Median, solid line. Mean, dashed line. Wilcoxon signed-rank test, p-value shown on plot.

We identified the z-position of each rearrangement initiation site where a high-order vertex first forms via contraction of a cell-cell contact, and we found a gradient in initiation frequency along the apical-basal axis (Fig. 2B, C). Half of the rearrangements analyzed initiated within the most apical 4 microns, but we also found significant numbers of rearrangements initiating all along the apical-basal axis (Fig. 2C). This was unexpected as we had anticipated that rearrangement initiation would be restricted to apical adherens junctions (8, 9) or basal sites of protrusive activity (21). Lateral rearrangement initiation has recently been observed in the neighboring presumptive mesoderm tissue, but is specifically associated with the presence of a second, more lateral tier of adherens junctions in those cells (41). We also identified a subset of multifocal rearrangements with nearly simultaneous initiation at both apical and more basolateral z-positions (Fig. 2B). We categorized rearrangements into four groups based on their initiation type (Fig. 2B). Initiation in the top 0-5 microns around the level of adherens junctions (apical) was most prevalent at 60±19%. Initiation in the next 10 microns (lateral) was next most prevalent at 27±17%. Initiation at z-positions below 15 microns (basolateral) or at multiple sites (multifocal) were relatively rare at 6±7% and 7±6% respectively (Fig. 2D). These results demonstrate that cell rearrangements initiate in a gradient along the apical-basal axis that is reminiscent of the gradients observed in 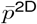 and *Q*, suggesting a link between tissue structure variations and cell rearrangement initiation along the apical-basal axis.

Next, we examined how cell rearrangements proceed after initiation by manually tracking the propagation of high-order vertex formation along the apical-basal axis over time (Fig. 2E, F). For all rearrangement types, we found that once initiated, the high-order vertex propagates along the apical-basal axis (Fig. 2F). Apical initiations propagate basally; basolateral initiations propagate apically; lateral initiations propagate both apically and basally; and multifocal initiations propagate laterally. We noticed that the velocity of vertex formation or “vertex velocity” seemed to vary as the vertex moved along the apical-basal axis (Fig. 2F, G). For roughly 70% of apically initiated rearrangements, vertex velocity was slower near the apical side of the tissue and increased as the vertex moved basally (Fig. 2G,H), similar a previous observation (41). The change in velocity often occurred at tissue depths between 5 and 10 microns, similar to the depths at which 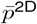 and *Q* decreased significantly (c.f. Fig. 1F’,G’). In contrast, for laterally initiated rearrangements, vertex propagation towards the apical side was faster than towards the basal side (Fig. 2I). Overall, we find that typical vertex propagation velocities are ≈ 5*µ*m/min. For comparison, this is faster than the ≈1 *µ*m/min velocity of furrow ingression during the fast phase of cellularization that forms the germband epithelium (42, 43). It is possible that vertex propagation speeds reflect some type of active self-propagating contractile activity or passive tissue mechanical response that varies with tissue depth.

Taken together, these results demonstrate that cell rearrangements initiate in a gradient along and then propagate at speeds linked to tissue depth across the apical-basal axis. Notably, the gradient in initiation frequency mirrors the gradient in tissue structure along the apical-basal axis, suggesting a link between high 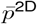 and rearrangement initiation rates. Moreover, the observed changes in vertex propagation velocities along the apical-basal axis suggest that the local structural environment of the tissue is linked with local cell contact remodeling.

### Complex 3D cell geometries are intermediate states during dynamic cell rearrangements

Our observations of high-order vertex formation followed by propagation along the apical-basal axis led us to investigate the 3D shapes of cells and their lateral cell contacts during cell rearrangements. We used single-cell tracking and a custom 3D reconstruction pipeline to represent the geometry of lateral cell-cell contacts as triangulated surfaces based on information from 2D cell outline networks in each z-plane (Fig. 3A-E, Materials and Methods).

**Fig. 3.**
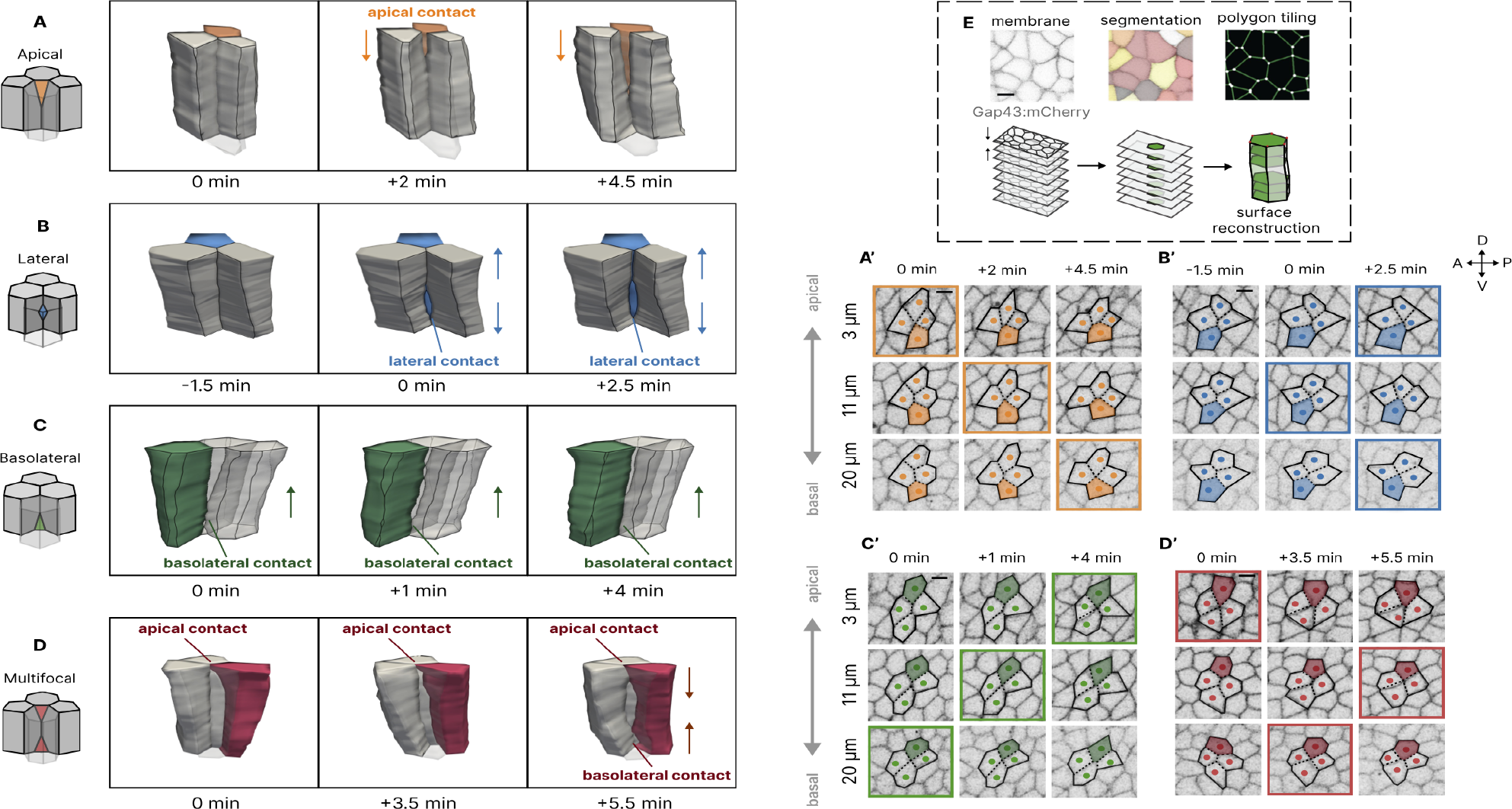
Complex 3D cell geometries are intermediate states during dynamic cell rearrangements. 3D reconstructions of rearrangements with apical **(A)**, lateral **(B)**, basolateral **(C)**, and multifocal **(D)** initiation, highlighting the variety of transitional cell geometries involved in dynamic cell rearrangements. Rearrangements with apical or basolateral initiation are associated with transient scutoids. Rearrangements with lateral or multifocal initiation are associated with more complex cell geometries and lateral cell-cell contacts. **(A’-D’)** Confocal fluorescence images of the example rearrangements from (A-D) at multiple z-planes and time points. In each panel, cell edges and contacts are highlighted with solid lines and dotted lines respectively and frames with high-order vertices are boxed. Scale bars, 5 microns. **(E)** Methods for 3D reconstruction of cell shapes and lateral cell-cell contacts as triangulated surfaces based on confocal imaging data.

We found a variety of complex 3D cell geometries that are transient intermediate states as cells change shape during rearrangements (Fig. 3A-D). These cell geometries correspond to distinct rearrangement initiation types: rearrangements initiated apically and basolaterally involve transient scutoids (Figs. 3A, A’, C, C’), while rearrangements initiated laterally or multifocally involve more complex cell contact patterns (Figs. 3B, B’, D, D’). These transient, intermediate cell geometries are typically resolved as the new lateral cell-cell contact first expands along the apical-basal axis to form a tall and narrow facet that subsequently widens; cells eventually recover their geometries as pyramidal frusta (Figs. 3A-D).

Scutoids were first identified as equilibrium cell shapes in curved epithelia (24, 29) and have recently been reported as transient states following cell proliferation in the sea star embryo (28). The transient scutoids we find here are distinct in that they occur in a non-proliferating epithelium in which cell numbers are not changing, so that the transient scutoids must be induced by other factors, such as dynamic actomyosin-generated stress patterns that vary along the apical-basal axis of the tissue. In the germband, these stress patterns likely arise from the planar polarized myosin pattern at apical adherens junctions and are linked with the observed gradients in tissue structure along the apical-basal axis. The basolateral rearrangements we find here may be related to rearrangements initiated by basal protrusive activity in the germband (21), although we do not resolve the most basal side of the tissue in our analyses. To our knowledge, the transient cell geometries we observe during lateral and multifocal rearrangements represent new classes of epithelial cell geometries.

### Lateral cell contact remodeling during rearrangement is a multi-step, multi-rate process

Remodeling of lateral cell-cell contacts is the fundamental process mediating epithelial cell rearrangements, but the nature of lateral contacts is poorly understood. This is due in large part to the challenge of quantitatively analyzing lateral cell surfaces compared to the more accessible apical or basal planar cross-sections in which cell contacts appear as polygon edges. To explore the process of lateral cell contact remodeling, we leveraged our reconstructions of lateral cell contact surfaces to quantify how existing contacts contract and new contacts grow during cell rearrangements (Fig. 4A-A’).

**Fig. 4.**
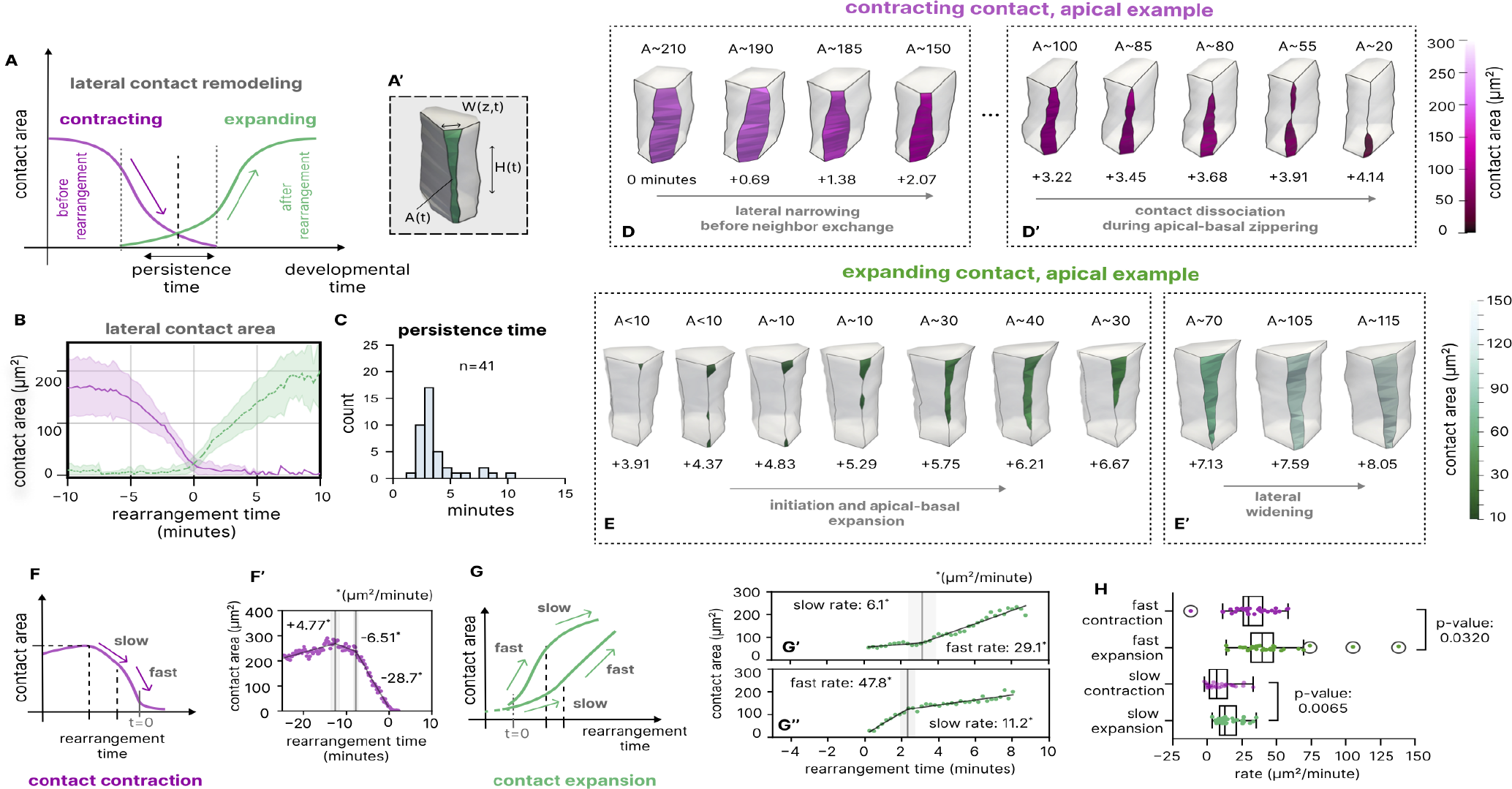
Lateral cell contact remodeling during cell rearrangement is a multi-step, multi-rate process. **(A)** Model of lateral cell contact remodeling during a cell rearrangement (T1 process), highlighting the time evolution of contracting and expanding lateral contacts; the persistence time is defined by the interval where both the contracting and expanding contacts are present (Materials and Methods). **(A’)** Schematic illustrating the apical-basal and lateral growth of a new contact. **(B)** Averaged time-series of areas of contracting and expanding contacts during cell rearrangement, centered by the rearrangement time *t* = 0 for n=41 rearrangements in N=2 embryos. **(C)** Distribution of persistence times. **(D’, D’)** Sample timelapse of a contracting contact in an apically initiated rearrangement; highlighted are two distinct steps in the process: initial lateral narrowing of the contact (D) followed by disappearance of the narrow contact along the apical-basal axis (D’); color indicates the value of the contact surface area. **(E, E’)** Sample timelapse of an expanding contact; highlighted are two distinct steps in the process: initial establishment of a narrow and tall contact (E) followed by widening of the contact (E’); color indicates the value of the contact surface area. **(F, F’)** Schematic and example contact area time-series showing a contracting contact with a rate change from slow to fast contraction; the three numbers in (F’) are the rates of pre-expansion, slow contraction, and fast contraction (microns sq./min). **(G, G’, G”)** Schematic and example contact area time-series illustrating the behavior of an expanding contact with slow then fast expansion (G’), and one with fast then slow expansion (G”). **(H)** Comparison between the slow and fast rates of contact contraction and expansion defined in (F-G”). The slow rates of contraction and expansion, as well as the fast rates of contraction and expansion, were statistically significantly different.

For each cell rearrangement, we measured the areas of the contracting and expanding contacts and defined *t* = 0 min as the time point at which these curves cross (Fig. 4A, B). On average, full contact expansion or contraction each take ∼ 5-10 min, and these two processes display temporal overlap (n=41 rearrangements in N=2 embryos) (Fig. 4B). We defined a persistence time for each rearrangement as the interval spanning the first appearance of the expanding contact and the last disappearance of the contracting contact (Fig. 4A). Equivalently, the persistence time is the lifetime of the transient, intermediate cell geometries during rearrangements (c.f. Fig. 3). We found that persistence times are broadly distributed with a mean of 3.75 ± 1.90 min (CV = 0.51; n=41 rearrangements in N=2 embryos) (Fig. 4C). These persistence times are longer than, but have a similar broad distribution as, those reported for 2D analyses of vertex lifetimes at the apical side of the germband (35, 44, 45) and 2D vertex model simulations of tissues with active junction remodeling (46). We did not find significant differences in persistence times between rearrangements initiated at different apical-basal positions.

Next, we analyzed the rates of lateral contact contraction and expansion over a time interval that included the first onset of contact contraction before neighbor exchange started and followed the growing contact after neighbor exchange concluded (Fig. 4A, D-E’). These rates are distinct from but related to vertex velocities (Fig. 2), as they capture changes in both contact height *H*(*t*) and width *W* (*z, t*) before and after neighbor exchange (Fig. 4A’). Overall, we found distinct rates of area change that are linked with steps in the rearrangement process (Fig. 4A, D-H). In the first step of a typical rearrangement, the existing lateral contact narrows (Fig. 4D). Next, the new lateral contact forms and expands vertically along the apical-basal axis into a tall, narrow facet, while the existing lateral contact contracts and disappears (Fig. 4D’, E). Finally, the new contact widens (Fig. 4E’).

Focusing first on contracting contacts, we find that the majority of analyzed contacts initially contract at a slow rate (9.4 ± 9.3 *µ*m^2^/min) before transitioning to a fast rate (33 ± 15 *µ*m^2^/min) (28/37 rearrangements in N=2 embryos) (Fig. 4F, F’). In the other rearrangements, we find a fast rate that is similar to the fast rate in the first group (30 ± 9 *µ*m^2^/min; 9/27 rearrangements in N=2 embryos) but with no discernible initial slow rate. We don’t find significant differences in these behaviors between rearrangements initiated at different apical-basal positions. The fast area contraction rate is consistent with the range of observed vertex propagation velocities (c.f. Fig. 2H, I), assuming the contact is a triangle with fixed base width and a height that decreases at the vertex propagation velocity. Comparing reconstructions of lateral contacts, we see that the initial slow rate corresponds to a narrowing of the contact, and the fast rate corresponds to rapid contraction or “zippering” of that tall, narrow facet as the newly formed high-order vertex propagates along the apical-basal axis (Fig. 4D,D’).

For the expanding contacts, we also find two rates of growth but with more variation in behaviors between rearrangements. In half of the analyzed rearrangements (17/34 rearrangements in N=2 embryos), we observe what appears to be an initial slow rate (14 ± 10 *µ*m^2^/min) and then a fast rate (42 ± 23 *µ*m^2^/min) (Fig. 4G’). In the other half (17/34 rearrangements in N=2 embryos), the contact grows at a fast rate before switching to a slow rate (Fig. 4G”). The fast and slow expansion rates were not significantly different between the two groups of expanding contacts. For expanding contacts, these rate changes appear to be linked with transitions between periods of apical-basal expansion and lateral widening (Fig. 4E,E’).

Taken together, these findings illuminate the nature of lateral cell contact remodeling during epithelial cell rearrangements in 3D. We find that lateral contact remodeling proceeds via a *geometric switch* between periods of lateral widening or narrowing and apical-basal contraction or expansion. This progression may reflect geometric or mechanical constraints on remodeling contacts in the germband epithelium. The broad distribution of persistence times and variation in lateral cell contact remodeling behaviors are signatures of the rich cell dynamics below the apical tissue surface that give rise to epithelial tissue remodeling during convergent extension.

## Conclusions

We have shown that cell rearrangements in the *Drosophila* germband initiate at positions along the apical-basal axis of the tissue in a pattern that matches a gradient in cell packing structure along this axis. The apical-basal gradients in 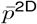 and *Q* that we find around the onset of GBE are consistent with the appearance of planar polarized patterns of actomyosin near apical adherens junctions that induce anisotropic patterns of stress at this side of the tissue (7, 8, 34– 36). The attenuation of structural changes towards the basal side of the tissue highlights the challenge of coordinating mechanical behaviors at the apical and basal sides of this tall epithelium. Although many cell rearrangements do initiate near the apical side of the tissue, our findings demonstrate that rearrangement initiation is not strictly localized to the apical domain, where contractile actomyosin is enriched (7–9), or to the basal domain, where protrusive activities have been reported (21). These findings raise the possibility that laterally initiated rearrangements are actively driven by lateral actomyosin contractility or are instead passive in nature (47).

In addition, we have shown that cell rearrangements involve the apical-basal propagation of high-order vertices and lateral cell contact remodeling. Vertex propagation velocities vary as vertices traverse the apical-basal axis and depend on the direction of travel, indicating that differing mechanical and molecular environments influence this process. Our pipeline for reconstructing and analyzing lateral cell-cell contacts allowed us to see that lateral contacts grow and shrink in a multi-step, multi-rate process, with new contacts first elongating along the apical-basal axis to form tall, narrow facets before widening. Notably, we find that the cell rearrangement process involves transient cell geometries, such as scutoids, that are distinct from previously reported stable scutoids in curved tissues (24) or transient scutoids in proliferating tissues (28), revealing the complex cell dynamics below the apical tissue surface during convergent extension.

These findings raise the central question of what mechanisms drive cell rearrangement along the apical-basal tissue axis. More specifically, are these processes actively driven by local cell-generated forces distributed all along the apical-basal axis or instead the result of active stresses exerted at the apical surface that then passively propagate basally due to tissue mechanics. The findings and approaches reported here will be a foundation for tackling this question. More broadly, the principles of 3D cell rearrangements we have uncovered and the approaches for quantifying 3D epithelial tissue structure we have developed should be broadly applicable to other epithelia that remodel during development or in disease.

## Materials and Methods

### Fly stocks and genetics

Embryos were generated at 23°C and analyzed at room temperature. Embryos were progeny of *yw* females with two copies of a sqh-gap43:mCherry transgene to label cell membranes (48).

### Live imaging

Embryos aged 2-4 hours were dechorionated for 2 min in 50% (vol/vol) bleach, washed in distilled water, and mounted in a 1:1 mixture of halocarbon oils 27 and 700 (Sigma) between a coverslip and an oxygen-permeable membrane (YSI). Embryos were oriented with the cephalic and ventral furrow visible at the edges of the field of view. A 212 µm x 159 µm ventrolateral region of the embryo was imaged *in vivo* on a Zeiss LSM880 laser-scanning confocal microscope using the 561 nm laser line, 40X/1.2 NA water-immersion objective, and the Zeiss AiryScan detector. Z-stacks were acquired at 1 micron steps and 13-15 second intervals with in-plane resolution of 0.33 microns x 0.33 microns. Mean intensity z-projections of 3 consecutive slices were analyzed at each z-position along the apical-basal axis.

### Image analysis

#### Apical z-position determination and z-drift correction

A kymograph was performed on a line parallel to the apical-basal axis that bisected the midsagittal view of the embryo to determine the location of the most apical z-plane of the tissue at all time points. If the location of the most apical z-plane varied over time, manual registration of z-planes was performed. The apical z-plane was defined as z=0 for multiple embryo comparison.

#### Measurement of tissue elongation along the anterior-posterior axis

Tissue elongation was measured by particle image velocimetry (PIV) using `openPIV` in `Python`. Each image was divided into 2-pass Fast-Fourier-Transform windows (50 × 50 pixels) with 30% overlaps. A displacement vector field for each window and each time point was determined by cross-correlating each window in the current time point and the image in the next time point. Tissue length change was measured by quantifying the cumulative sum of the anterior-directed displacement at the anterior end of the germband and the posterior-directed displacement at the posterior end of the germband. The onset of tissue elongation (*t* = 0) was defined as the time point when the derivative of the tissue elongation curve intersects zero at the most apical side of the tissue. Embryos were registered in time using the onset of tissue elongation for multiple embryo comparisons.

#### Cell segmentation and tracking

We segmented 3D movies of gap43:mCherry one z-plane at a time, restricting analysis to z-planes with sufficient image quality for segmentation (typically z=2-21 microns). First, we generated automated cell segmentations for each z-plane using a fine-tuned instance of the `cyto2` model in `CellPose 2.0` (49). Next, we processed each z-plane using morphological operations in `scikit-image` (50) to remove holes, isolated connected components, and any spaces between cell masks until a confluence criterion was met. We then tracked cells over each z-plane using `btrack` (51) and coalesced the tracks using the `stitch3D` function in `CellPose`. We used these data in 3D reconstruction of lateral cell-cell contacts. Finally, we manually inspected select z-planes and corrected the segmentations as needed in the `MATLAB` package `SEGGA` (36). Selected z-planes for manual correction consisted of the most apical and basolateral z-planes and subsequent z-planes in between at intervals of 2-5 microns. `SEGGA` (36) was also used to track cells over time in pre-defined regions of interest in each z-plane. These data formed the basis for the analysis of cell and tissue morphology.

#### Identification and tracking of cell rearrangements

For the analysis reported in Fig. 2, we identified a subset of all cell rearrangements in each movie by visual inspection and manual tracking of the positions of the associated high-order vertices over time and over the apical-basal axis as shown in Figs. 2 and 3. Cell rearrangements used for analysis were identified by searching for high-order vertices that formed from a shrinking edge and resolved into a new edge at all z-positions within the ROI during the movie. Classification of the identified cell rearrangement was based on the initiation location falling within the set ranges along the apical-basal axis (Fig. 2D). If consecutive z-positions initiated at the same time, the median z-position was used as the initiation location. Higher-order vertex formation and resolution into edges were tracked for identified cell rearrangements. Only cell rearrangements involving four cells (T1 processes) were included in analyses.

#### High-order vertex velocity analysis

For unidirectional rearrangements (apical and basolateral), velocity was calculated by dividing the distance traversed (18*µ*m) by the difference in time between high-order vertex formation at the apical most z-position and basal most z-position. For multi-directional rearrangements (lateral and multifocal), two velocities were calculated by dividing the distance between initiation (lateral) or termination (multifocal) position and apical or basal most z-position by the difference in time between higher-order vertex formation at initiation or termination position and apical or basal most z-position. Higher-order vertex propagation velocity for multi-directional rearrangements was divided into apical or basal direction of propagation. Velocity data for each rearrangement type was combined from three embryos for statistical analysis as the difference between the average propagation velocities in the three embryos was not statistically significant. Velocity data for basolateral and multifocal rearrangements were automatically combined from three embryos due to small sample size.

#### Velocity breakpoint analysis

We used piece-wise (segmented) linear regression (52) of vertex propagation along the apical-basal axis to estimate potential changes in vertex propagation velocity. We allowed the preset number of break-points to vary between 0 and 3 and visually inspected each fit to select the best fits. We used the `Python` package `piecewise-regression` (53).

#### 2D tissue network reconstruction

Analysis of the cell outline network topology and 3D reconstructions required us to convert the cell segmentations into planar tilings with explicitly defined nodes and edges that formed each polygonal cell. We extracted the cell outline network in each z-plane from cell outlines skeleton masks and converted the skeleton mask into a graph where multicellular junctions formed the nodes and bicellular junctions formed the edges, and each cell was represented as a polygon. These data formed the basis for the calculation of the vertex coordination number Ƶ, the cell shape index *p*^2*D*^, the triangle elongation tensor *q*, the cell shape alignment index *Q*, and the cell rearrangement rates.

#### 3D cell and tissue reconstruction

We used a custom 3D reconstruction pipeline to infer 3D cell geometry from segmented z-stacks of the cell outline marker. To ensure a confluent tissue and spatially consistent reconstructions, we used a constrained 3D cell surface triangulation procedure as follows: (i) a cell outline skeleton mask was extracted from the cell segmentations in each z-plane; (ii) the skeleton mask was converted into a graph by identifying and labeling the nodes and edges of the cell outline network; (iii) the set of nodes and edges that defined each lateral cell-cell contact were identified and isolated across the entire z-stack; (iv) lateral cell-cell contact surfaces were triangulated while using the constituent nodes and edges as spatial constraints; (v) finally, each cell was reassembled as a set of lateral, apical, and basal triangulated surface patches. This approach avoids the common pitfalls of conventional surface triangulation recipes like the marching cubes algorithm and preserves the curvature distributions of lateral cell-cell contact surfaces when applied to data with anisotropic voxel resolutions. It also provides direct access to the morphology and deformations of each individual cell-cell contact. We built the pipeline in python using `scikit-image` (50), `sknw` (54), and `pyvista` (55). We used ParaView (56) to create the 3D renderings in Figs. 3 and 4.

### Quantification of cell and tissue structure

#### Vertex coordination number

Each vertex of the cell contact network has an associated coordination number Ƶ, equal to the number of edges it connects. The ROI-averaged vertex coordination number 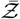 is the average number of cells meeting at a single vertex. A threshold edge length of 0.75 microns was determined manually and led to results consistent with prior reports (19).

#### Cell rearrangement rate

We measured the cell rearrangement rate in each z-plane by tracking the number of neighbors lost per cell per unit time in the ROI using `SEGGA` as described in (36).

#### Cell shape index

The 2D cell shape index is defined as the ratio of the cell perimeter to the square root of cell area, 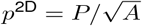 (Fig. 1). The tissue average cell shape index, 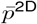, is the mean of *p*^2D^ over all cells in a given region of interest. We computed the cell shape index for polygon cell outlines unless stated otherwise.

#### Cell shape alignment index

The cell shape alignment index was computed using a triangulation of the cell outline network as described in (31) and demonstrated in (19). Briefly, the tissue plane is tiled with triangles that are formed by connecting the cell barycenters and high-order vertices (those with Ƶ *>* 3) (Fig. 1). In the resulting tiling, the elongation, *q*, of each triangle with respect to an equilateral reference triangle provides a measure of local cell shape anisotropy and the area-weighted average ***Q*** = ⟨***q***⟩ provides an aggregate measure of shape anisotropy of the cell network. The magnitude of this tensor, 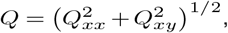, called the *cell shape alignment index*, provides a scalar measure of tissue anisotropy. The triangle elongation tensor, *q*, and the cell shape alignment index, *Q*, were both calculated using the python package `triangles-segga` (19).

#### Analysis of lateral contact remodeling during rearrangements

We analyzed the geometry and kinetics of lateral cell-cell contact remodeling by 3D reconstruction of contact surfaces to measure the change in contact surface areas over time. Each pairwise contact surface was triangulated as described previously, and we calculated the surface area by summing up the areas of the constituent triangles.

#### Rearrangement time

We estimated a *rearrangement time* defined as the cross-over point of the contracting and expanding contact area time-series (Fig. 4A). We first identified the contracting and expanding contact surfaces in each group of four cells involved in a rearrangement. We then found the point where the two curves intersected after smoothing both curves using a boxcar moving average filter with size equal to 20 time-points for the calculations reported in Fig.4.

#### Persistence time

We defined the *persistence time* as the interval defined by (i) the growing edge increasing beyond a minimum threshold surface area on one hand, and (ii) the contracting edge surface area decreasing below the same minimum threshold on the other. We used a minimum threshold surface area of 10 µm^2^ in the calculations reported in Fig. 4.

#### Contact remodeling rates

We used piece-wise (segmented) linear regression (52) of the contact area time-series to estimate the rates of contact contraction and expansion during rearrangements and potential rate changes in each curve. We allowed the preset number of breakpoints to vary between 1 and 3 and visually inspected each fit to select the best fits. We used the `Python` package `piecewise-regression` (53). We used the Mann-Whitney U test to compare the rates of contact contraction and expansion for the results reported in Fig. 4.

## Data Availability

Data will be available by request from the corresponding author.

## Software Availability

The pipeline for 3D reconstruction of lateral cell-cell contacts will be made publicly available.

## ACKNOWLEDGEMENTS

We thank members of the Kasza Lab for helpful comments on the manuscript. This work was supported by NIH Grant R35GM138380 to KEK; KEK holds an NSF CAREER Award, Packard Fellowship, and Sloan Research Fellowship in Physics.

## AUTHOR CONTRIBUTIONS

XW and KEK conceived the initial studies of tissue structure variation along the apical-basal axis. XW performed proof-of-concept experiments and initial analyses of tissue structure. EMK and KEK conceived the manual analyses of vertex dynamics and cell rearrangements along the apical-basal axis. SO and KEK conceived the analyses of lateral cell-cell contact remodeling. EMK generated the fly stocks and performed the live imaging experiments. SO and EMK developed the image analysis pipeline. EMK performed manual analyses of high-order vertex initiation and propagation. SO and XL designed the 3D reconstruction pipeline. XL implemented and tested the 3D reconstruction pipeline. SO, EMK, and XL performed the quantitative analyses of cell-level and tissue-level live imaging data. SO, EMK, XW, XL, and KEK wrote the manuscript.

